# ifCNV: a novel isolation-forest-based package to detect copy number variations from various NGS datasets

**DOI:** 10.1101/2022.01.03.474771

**Authors:** Simon Cabello-Aguilar, Julie A. Vendrell, Charles Van Goethem, Mehdi Brousse, Catherine Gozé, Laurent Frantz, Jérôme Solassol

## Abstract

Copy number variations (CNVs) are an essential component of genetic variation distributed across large parts of the human genome. CNV detection from next-generation sequencing data and artificial intelligence algorithms has progressed in recent years. However, only a few tools have taken advantage of machine learning algorithms for CNV detection, and none propose using artificial intelligence to automatically detect probable CNV-positive samples. Furthermore, in general, most CNV software that is developed for specific data types has sub-optimal reliability for routine practice. In addition, the most developed approach is to use a reference or normal dataset to compare with the samples of interest, and it is well known that selecting appropriate normal samples represents a challenging task which dramatically influences the precision of results in all CNV-detecting tools. With careful consideration of these issues, we propose here ifCNV, a new software based on isolation forests that creates its own reference, available in R and python with customisable parameters. ifCNV combines artificial intelligence using two isolation forests and a comprehensive scoring method to faithfully detect CNVs among various samples. It was validated using datasets from diverse origins (capture and amplicon, germline and somatic), and it exhibits high sensitivity, specificity and accuracy. ifCNV is a publicly available open-source software that allows the detection of CNVs in many clinical situations.

**Key points:**

- Copy number variation detection
- Machine learning
- Localisation scoring
- Benchmark on various clinical situations and on various datasets
- Easy-to-use R and Python open-source Package

## Introduction

Copy number variations (CNVs) are a class of structural variations that result from the deletion or duplication of a DNA fragment. About 1500 CNV regions have already been discovered in the human population, accounting for ~12–16% of the entire human genome,^1^ making it one of most common types of genetic variation. Although the biological impact of the majority of these CNVs remains uncertain, nearly 50% of known CNVs overlap with protein-coding regions, and many are involved in genetic diseases. Recent studies have demonstrated that CNVs can be implicated in many rare diseases, such as inherited retinal dystrophies,^2^ and in diseases that involve dosage-sensitive developmental genes, such as Charcot-Marie-Tooth disease^3^ and DiGeorge syndrome among others.^4–6^ CNVs, resulting from gene amplification (copy number gain) as well as gene deletion (copy number loss), are common in cancer cells, and multiple studies have shown that duplication or deletion of specific genes can contribute to tumour growth^7^ and to resistance to anti-tumour therapies.^8,9^ In cancer cells, the size of these molecular alterations can vary dramatically, from one or a few exons to an entire chromosomal arm.

Although most CNVs found in cancer cells are likely to have accumulated, as a direct consequence of clonal evolution during the disease course, some have been identified as playing a role in the early development of cancer (e.g. CNVs located in *BRCA1/2* in familial breast and ovarian cancer^10^).

In fact, it has been estimated that CNVs represent more than 10% of the molecular alterations linked to cancer predisposition, making their detection a priority. Detection of acquired (somatic) focal copy number changes is also required for diagnosis, prognosis, and for the therapeutic management of patients with cancer.^11^ For example, loss of chromosomal arms 1p and 19q is closely associated with oligodendrogliomas, a subtype of primary brain tumours, and with a favourable prognosis in diffuse gliomas.^12^ Focal copy number increases are biomarkers predictive of responses to particular therapies; for example, patients with oncogenic *ERBB2* amplification in breast cancer respond well to trastuzumab, and acquired resistance to tyrosine kinase inhibitors is exhibited in patients with *MET*-amplified non-small cell lung carcinomas.^13,14^

Despite their importance in cancer and other diseases, detecting CNVs has remained challenging. CNV detection in clinical laboratories has traditionally relied on techniques such as karyotyping, multiplex ligation and probe amplification (MLPA) and fluorescence *in situ* hybridisation. These techniques, however, are often low-throughput and labour intensive. Over recent years, diagnostic laboratories have established higher-throughput methods, based on arrays such as comparative genomic hybridisation (aCGH) and single nucleotide polymorphism arrays,^15^ which can type thousands of known variants in the genome. Although these techniques are still routinely used, the rapid implementation of high-throughput next-generation sequencing (NGS) methods, especially targeted DNA panels, in clinical laboratories has led to the emergence of a fairly large number of pipelines and algorithms able to detect CNVs from NGS data.^16–28^

Most of these studies use the read-depth approach, relying on the hypothesis that the number of reads aligned to a genomic region is proportional to the copy number of the region. In multiple sample methods, CNVs are detected by comparing the read counts of the sample of interest to the read counts of a reference sample. The proper building of the reference is one of the main difficulties. To that end, two main solutions exist: (i) to gather a database of normal samples or (ii) to add normal samples into the NGS run. Nevertheless, both solutions come with issues, mainly the presence of a batch effect and a high cost, respectively. To avoid these problems, use of the single sample method was previously proposed, which consists of statistical modelling of the target read counts within the sample of interest to detect CNVs. Recent advances in artificial intelligence and in particular the availability of accessible machine learning packages^29^ has made it possible for developers to improve their algorithms in many areas. To date, only a few studies have taken advantage of these recent developments in the field of CNV detection from NGS data.^16,17,27^ An interesting example is CNV_IFTV^27^; in this study, the authors propose a single sample method in which the isolation forest (IF) algorithm^30^ is used to create an anomaly score and therefore detect CNVs. However, by the authors’ own admission, the algorithm runs for a considerable length of time, and they themselves turn towards a multiple sample method.^19^

With careful consideration of these issues, we present ifCNV, a novel machine learning-based software, provided as a python and R package (https://github.com/SimCab-CHU/ifCNV and https://github.com/SimCab-CHU/ifCNVR, respectively). It uses two IFs, one to split the CNV-positive (CNV^pos^) samples from the CNV-negative (CNV^neg^) ones, and a second to detect the altered targets within the CNV^pos^ samples. This approach combines several advantages, among which is that it does not require an external nor internal reference and it is low resource-consuming.

To validate our model and explore its limitations in clinical practice, we tested ifCNV on different synthetic datasets mimicking relevant clinical situations and on datasets obtained from amplicon- or capture-based DNA library preparation technologies.

## Materials and methods

### Isolation forest algorithm

The IF algorithm was developed by Liu et al.^30^ It ‘isolates’ observations using a binary tree structure called an isolation tree. In this isolation tree, anomalies are more likely to be isolated closer to the root, whereas normal points are more likely to be isolated at the deeper end. The IF algorithm builds its isolation trees for a given dataset by randomly selecting a feature and then randomly selecting a split value between the maximum and minimum values of the selected feature. The number of splittings required to isolate a sample is equivalent to the path length from the root node to the terminating node. This path length is then averaged over a forest of such isolation trees to produce a decision value. The smaller the value, the more likely it is that the sample represents an anomaly.

### Synthetic datasets

To create synthetic datasets that reproduce faithfully those obtained in routine clinical practice, we selected 1910 samples from in-house targeted NGS data with no CNVs. We extracted the total reads on each target from the aligned .bam files with the bedtools^31^ multicov function and created a reference reads matrix ordered with samples as columns and targets as rows. This reference reads matrix was then normalised by dividing each column by its median. All medians were used to create a median reads distribution that was needed for the rescaling process. Next, we created a normalised target reads distribution from each row of the normalised matrix. Thus, to generate synthetic datasets, we filled each line by taking a normalised target reads distribution, in which we randomly picked a value for each column. To rescale this matrix, we multiplied each column with a value randomly picked from the median distribution. Finally, to create CNV^pos^ samples within this synthetic dataset and test the algorithm, we modified the desired number of targets by multiplying it by a factor ranging from 0 to 10.

### ICR96 dataset

The dataset ICR96 exon CNV validation series^32^ was downloaded from the European Genome-phenome Archive (EGA) (EGAD00001003335). This dataset consists of the read counts of 96 samples sequenced on a targeted panel for which the copy number, at the exon level, was validated using high-resolution MLPA experiments (Table 1).

**Table 1.** Description of the datasets used in the study.

| Datasets | Samples | Positives | Negatives | Number of alterations | Amplifications | Deletions |
| --- | --- | --- | --- | --- | --- | --- |
| TSCA dataset | 81 | 25 | 56 | 39 | 21 | 17 |
| Juno Dataset | 43 | 26 | 17 | 28 | 9 | 19 |
| ICR96 dataset | 96 | 68 | 28 | 296 | 80 | 216 |

### In-house datasets

DNA extracted from clinical samples was analysed alongside by two molecular approaches: aCGH as a reference method and NGS using two different library preparation protocols.

#### DNA extraction of formalin-fixed paraffin-embedded samples

DNA was extracted from tissue samples using the Maxwell^®^ RSC DNA FFPE Kit (Promega, Madison, WI, USA) according to the manufacturer’s recommendations. DNA was quantified using the Qubit dsDNA BR Assay Kit and a Qubit Fluorometer (Thermo Scientific, Wilmington, DE, USA).

#### Comparative genomic hybridisation

aCGH profiling was performed with the Human Agilent Sureprint G3 8×60 K Microarray Kit (Agilent Technologies, Santa Clara, CA, USA). Tumour DNA was labelled with cyanine 5 (Cy5), while reference DNA from an individual of the same sex as the patient was labelled with cyanine 3 (Cy3). Sample and reference DNAs were pooled and hybridised for 24 hours at 67°C on the arrays. The fluorescence was read by an Agilent SureScan Microarray scanner and the Cy5/Cy3 fluorescence ratios were converted into log2 transformed values with Cytogenomics software (Agilent).

The threshold of the absolute value of the log2 fluorescence ratio retained to define a chromosomal anomaly was 0.25. A mean log2 ratio was calculated when, for at least three probes located on contiguous positions on the chromosome, a log2 ratio absolute value greater than 0.25 and of the same sign was measured. The minimum size of the anomalies considered in the interpretation of the results was set at one megabase (Mb).

The different chromosomal anomalies were defined by the Cytogenomics software according to the mean log2 ratio values, as follows: homozygous deletion for a value < −1, loss of one gene copy for a value between −0.25 and −1, gain of one gene copy for a value between 0.25 and 1 and amplification (gain of at least five copies) for a value > 2.

#### TruSeq Custom Amplicon library preparation assay – TSCA dataset

Library preparation was performed as previously described.^33^ Briefly, extracted DNA was qualified using KAPA Sybr^®^ Fast qPCR (Kapa Biosystems, Boston, MA, USA). A home-made panel targeting specific exons of 35 clinically relevant cancer genes was used for amplification of regions of interest. For each sample, dual-strand libraries were prepared using a TruSeq Custom Amplicon protocol, as described by the manufacturer (Illumina, San Diego, CA, USA). After amplification, PCR products were purified using AMPure XP beads (Beckman Coulter, Brea, CA, USA) and quantified, normalised and pair-end sequenced on a MiSeq instrument (2×150 cycles, Illumina). This dataset is composed of 81 samples from 59 different sequencing runs, with 25 CNV^pos^ samples (Table 1).

#### Advanta Solid Tumor NGS library preparation assay – Juno dataset

Libraries were prepared using the Advanta Solid Tumor NGS Library Prep Assay with the automated Juno™ system on integrated fluidic circuits (LP 8.8.6 IFC) (Fluidigm, San Francisco, CA, USA) following the manufacturer’s procedure. The panel is developed to allow the detection of somatic mutations in 53 oncology-relevant genes (234 kb, 1508 assays). Briefly, the LP 8.8.6 IFCs were primed with 20 ng of DNA per sample in the PCR mix. After amplification, pooled harvested samples were purified using AMPure XP beads (Beckman Coulter, Brea, CA, USA), and a second PCR was performed to integrate the sequencing adapters. Libraries were then quantified, normalised and pair-end sequenced on a NextSeq instrument (2×150 cycles, Illumina). In this dataset, there are 43 samples with 26 CNV^pos^ from 20 different sequencing runs (Table 1).

### Binary classification indicators

True positive rate (TPR or sensitivity), false positive rate (FPR), true negative rate (TNR or specificity), false negative rate (FNR), positive predictive value (PPV), accuracy (Acc) and the Matthews correlation coefficient (MCC) were used to measure the performance of ifCNV. These were computed as:

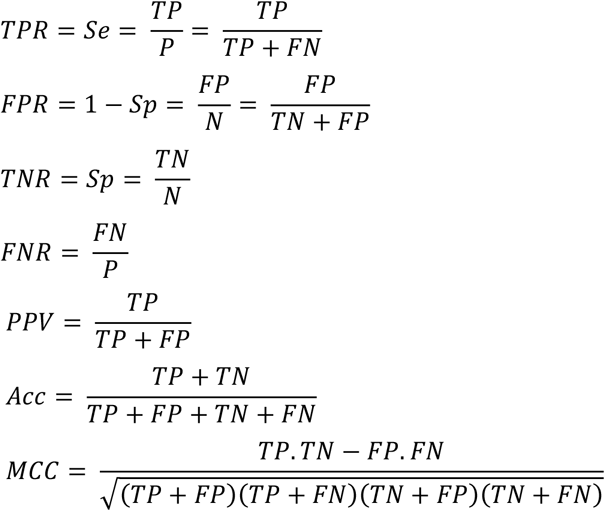

### Localisation scoring

Specific regions of biological significance (gene or exon) can be covered by several targets. In the event that a region is altered, all the targets in the region should be modified. By contrast, if only one target in the region is modified, it is likely to be an FP. We integrated this reasoning to develop a localisation score in order to reduce the number of FPs. The localisation score depends on the number of modified targets in the region, the number of targets in the region and the total number of targets in the panel. A semi-open log scale incorporating the ratio of modified targets in the region was chosen. It is calculated as follows:

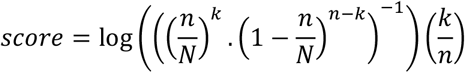

*with k the number of modified targets on the region*,
  *n the number of targets on the region*,
  *N the total number of targets*

### Pre-processing

A pre-processing step using the bedtools^31^ multicov function to generate the reads matrix was included in the package. It takes as input the aligned .bam files and outputs a read-depth matrix that was used for the CNV detection analysis. The alignment of the .fastq files on the reference genome (i.e. the creation of the .bam files) is not provided as a function in the ifCNV package and should be done by the user.

## Results

### ifCNV workflow

ifCNV is a CNV detection tool based on read-depth distribution obtained from NGS data (figure 1A). It integrates a pre-processing step to create a read-depth matrix using as input the aligned .bam files and a corresponding .bed file. This reads matrix is composed of the samples as columns and the targets as rows. Next, it uses an IF machine learning algorithm to detect the samples with a strong bias between the 99^th^ percentile and the mean (for amplifications, figure 1B, top plot), and the 1^st^ percentile and the mean (for deletions, figure 1B, bottom plot). These samples are assumed to be CNV^pos^. The samples with no bias, and therefore not detected by the IF as outliers, are considered as CNV^neg^ samples. After filtering of the samples with a mean read depth per target less than N (N=100 by default but can be set by the user to any value), the reads matrix is normalized by dividing each column (i.e., the reads distribution of each sample) by its median. Then, ifCNV creates a mean normalized normal sample by averaging all the CNV^neg^ samples. It is used as a reference to detect the outlying targets in each normalized CNV^pos^ sample with a second IF (figure 1C). These assumed altered targets, are then used to compute the localization score per region of interest (see methods section). Finally, a threshold is applied on this score to select the significantly altered regions that are compiled in an html report containing a table and a graph for easy user interpretation.

**Figure 1.**
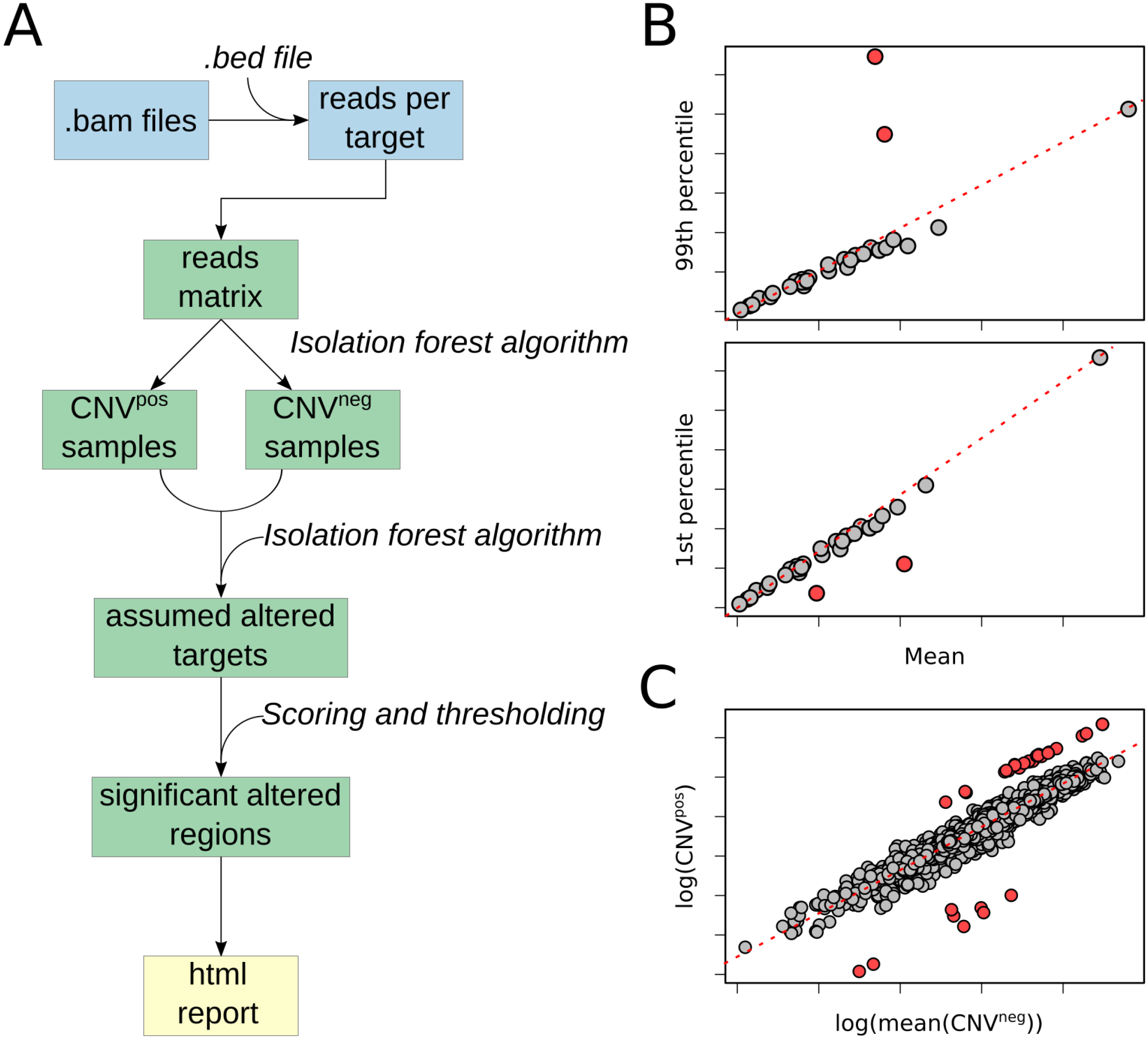
ifCNV workflow. A. ifCNV is composed of three steps: the pre-processing (blue), the core algorithm (green) and the output (yellow). CNV: copy number variation; CNV^neg^: CNV-negative samples; CNV^pos^: CNV-positive samples. B. top: 99^th^ percentiles of the reads distribution according to the means of the reads distribution of the samples in a NGS sequencing run; bottom 1^st^ percentiles of the reads distribution according to the means of the reads distribution of the samples in a NGS sequencing run. The red dots correspond to the outlying samples. C. In one CNV^pos^ sample, the logarithm of the reads per target according to the logarithm of the mean normalized normal sample. The red dots correspond to the outlying targets.

### Performance of ifCNV

#### Detection of CNV^pos^ samples

To quantify the ability of ifCNV to detect CNV^pos^ samples, we created a synthetic dataset of 1500 targets and 30 samples, in which we inserted one CNV^pos^ sample. It is noteworthy that if the copy number ratio (CNR) or the modified target ratio (MTR) are low, the CNV^pos^ samples will resemble the CNV^neg^ samples and therefore will be difficult to detect. Thus, the performance of a CNV detector directly depends on the CNR and the MTR. Taking this fact into consideration, we iterated the CNR and MTR (from 0 to 10 and 0 to 0.1, respectively) and performed 1000 simulations for each iteration (figure 2 A, B). The analysis of the attribution of CNV^pos^ samples for a CNR greater than 1 is shown in figure 2A. For CNRs greater than 6, ifCNV correctly identified the abnormal sample in 99.58% of simulations when the MTR is greater than 0.01 (figure 2A). Furthermore, if the CNR was between 4 and 6 and the MTR was over 0.01, ifCNV found the abnormal sample in 99.47% of simulations. Finally, if the CNR was over 2 and the MTR over 0.01, ifCNV detected the abnormal samples in 99.34% of simulations; this reached 99.83% when the MTR was greater than 0.035.

**Figure 2.**
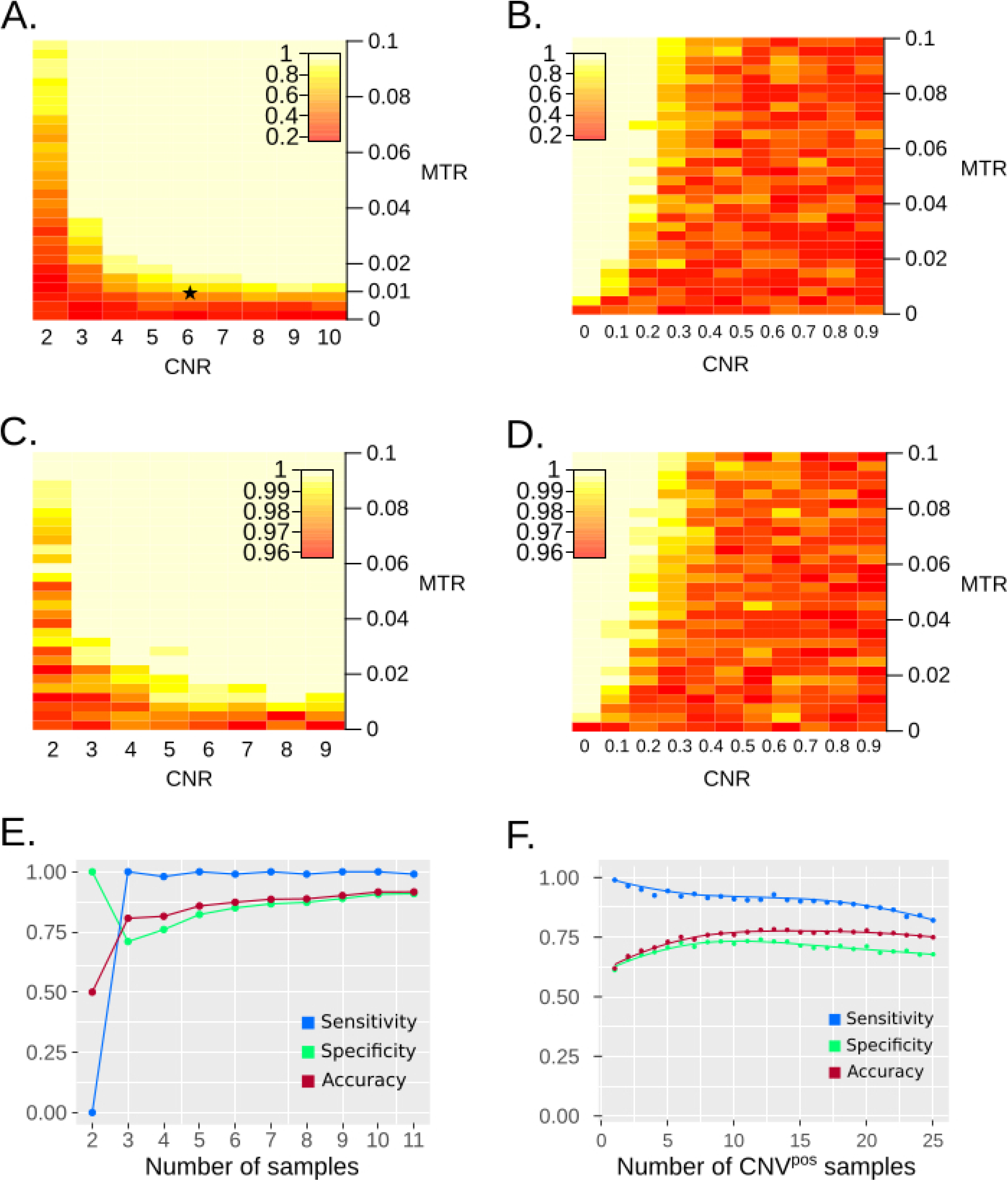
Performance assessment in detecting CNV^pos^ and CNV^neg^ samples. A. Heatmap of the detection rate of CNV^pos^ samples as a function of the CNR and the MTR, for CNRs between 2 and 10. B. Heatmap of the detection rate of CNV^pos^ samples as a function of the CNR and the MTR, for CNRs between 0 and 0.9. C. Heatmap of the detection rate of CNV^neg^ samples as a function of the CNR and the MTR, for CNRs between 2 and 9. D. Heatmap of the detection rate of CNV^neg^ samples as a function of the CNR and the MTR, for CNRs between 0 and 0.9. E. Classification indicators of the detection of a single CNV^pos^ sample in a set of several CNV^neg^ samples. F. Classification indicators of the detection of multiple CNV^pos^ samples in a set of several CNV^neg^ samples. CNR: copy number ratio; MTR: modified target ratio.

ifCNV was also able to detect deletion (CNRs under 0); as for CNRs greater than 1, the sensitivity was related to both the CNR and the MTR (figure 2B). For CNRs under 0.5, ifCNV detected the abnormal sample in 92.26% of simulations. For CNRs over 0.5, ifCNV only detected the abnormal sample in 27% of simulations. Although 27% is a higher detection rate than a random choice, for which the probability is 3.33% (1 sample out of 30), it is not satisfactory enough for diagnostic purposes given the importance of heterozygous deletion (hDel) in several diseases.^6^ To overcome this pitfall, we assessed an alternative solution based on the ability of the IF to accurately detect the CNV^neg^ samples. Indeed, samples labelled as CNV^neg^ were considered to be normal and were adopted as an internal reference.

#### Identification of the CNV^neg^ samples

To quantify the ability of ifCNV to detect CNV^neg^ samples, we used the same synthetic dataset, and we also iterated the CNR and MTR (from 0 to 10 and 0 to 0.1, respectively) and performed 1000 simulations for each iteration (figure 2 C, D). ifCNV was able to correctly identify the CNV^neg^ samples in 99.87% of simulations, regardless of the CNR and MTR. For CNRs over 2, this reached 99.9%. Interestingly, for CNRs under one, ifCNV identified the CNV^neg^ samples in 98.57% of the simulations. Finally, a value of 99.69% was obtained for CNRs under 0.5.

#### Detection of one CNV^pos^ sample in a set of several CNV^neg^ samples

Conventional hospital and research laboratories must determine the CNV status of numerous samples. The number of samples in a sequencing run can vary from a few to several dozen. We therefore assessed the ability of ifCNV to correctly find a unique CNV^pos^ sample in a set of several negative samples. We created synthetic datasets of 2 to 100 samples in which we inserted a CNV^pos^ sample with a CNR of 5 and an MTR of 0.03. We performed 100 simulations for each and calculated the sensitivities (Se), specificities (Sp) and accuracies (Acc) of the algorithm (figure 2E). For one CNV^pos^ sample in a set of two, ifCNV failed to label any sample as positive, leading to one TN and one FN (Se=0, Sp=1 and Acc=0.5). Interestingly, for one CNV^pos^ sample in three, ifCNV correctly labelled the positive sample in every simulation and found one FP in less than 50% of simulations. When increasing the number of samples in the set, the Se remained close to 1 (0.992 from 3 to 100 samples), and the Sp and the Acc tended to 0.9.

#### Detection of multiple CNV^pos^ samples in a set of several CNV^neg^ samples

As several CNV^pos^ can be present in the same sequencing run, we tested the performance of ifCNV in such situations. We randomly chose 2 to 25 samples in a synthetic dataset of 50 samples. We then added random CNRs (from 2 to 6) to 3% of the targets of these samples and performed 100 iterations to determine the Se, Sp and Acc of our algorithm (figure 2F). ifCNV exhibited relatively high Se, Sp and Acc (around 0.85, 0.7 and 0.75, respectively), regardless of the number of CNV^pos^ samples in the dataset. Notably, when half of the tested set (25/50) was CNV^pos^, ifCNV correctly labelled a mean of 20 samples across all simulations.

#### Detection of altered targets

The second step of ifCNV consists of labelling the targets that are modified among the CNV^pos^ samples. To assess its performance, we created a synthetic dataset of 30 samples and 300 targets. One sample was CNV^pos^ with one modified target randomly chosen at each iteration, with a CNR from 0 to 5. We then performed 1000 iterations and calculated the Se, Acc, PPV and MCC (figure 3). On the one hand, ifCNV exhibited a Se very close to 1 for CNRs from both 0 to 0.3 and 3 to 5, meaning that the modified target was accurately labelled in almost every simulation. On the other hand, the PPV and the MCC were low (~0.02 and ~0.1, respectively), reflecting a high number of FPs. However, the Acc was ~0.87 and stayed approximately unchanged, meaning that the number of TNs was high and dominated the number of FPs.

**Figure 3.**
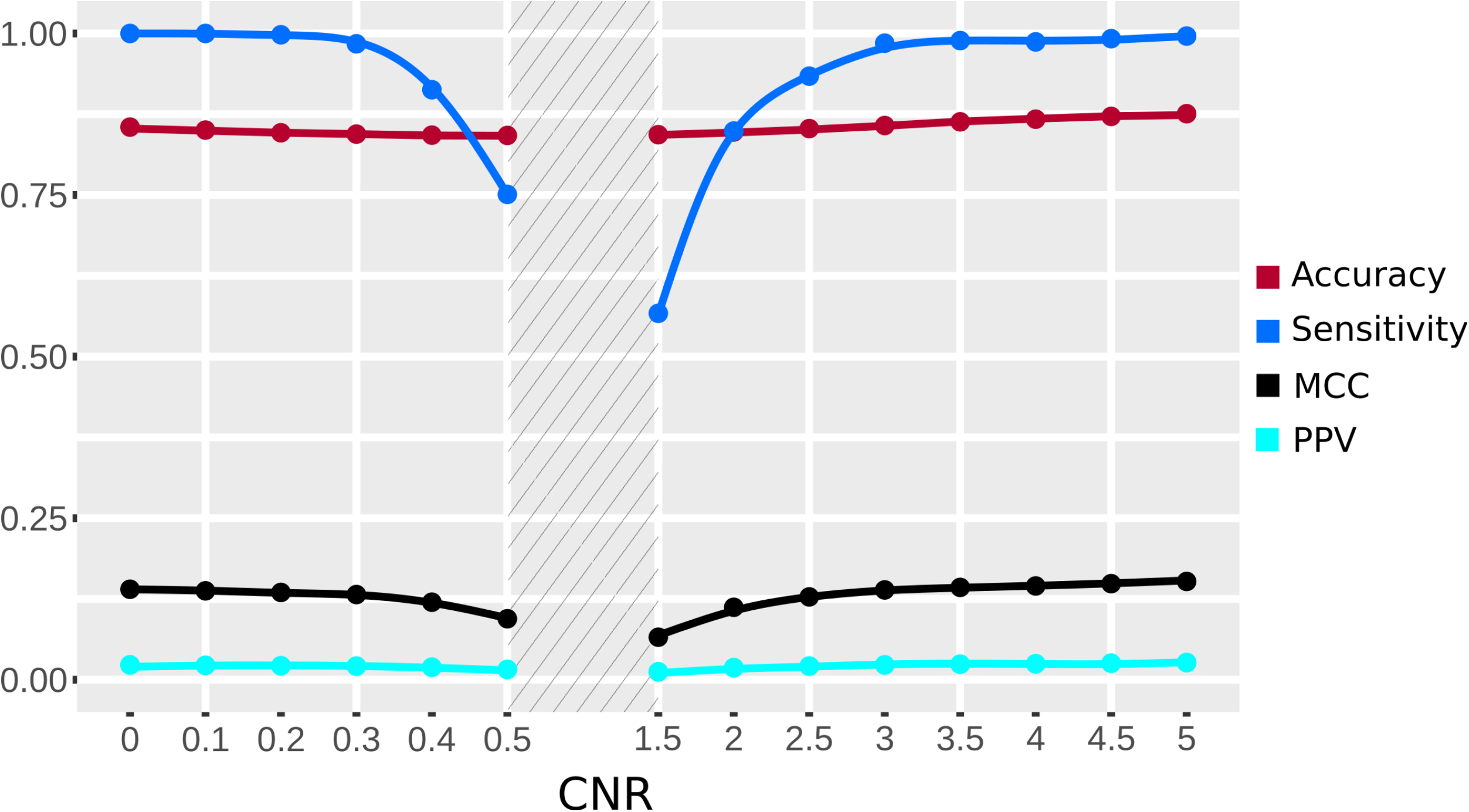
Classification indicators for the detection of altered targets. MCC: Matthews correlation coefficient; PPV: positive predictive value.

#### Thresholding on the localisation score

To test the ability of the score to discriminate the FPs from the TPs, we used the same synthetic dataset as before and grouped targets (from 2 to 15) together to mimic regions. We then modified all the targets corresponding to a randomly chosen region. Finally, we computed the localisation score and calculated the Se, Acc, PPV and MCC before (figure 4A, left panel) and after thresholding (figure 4A, right panel).

**Figure 4.**
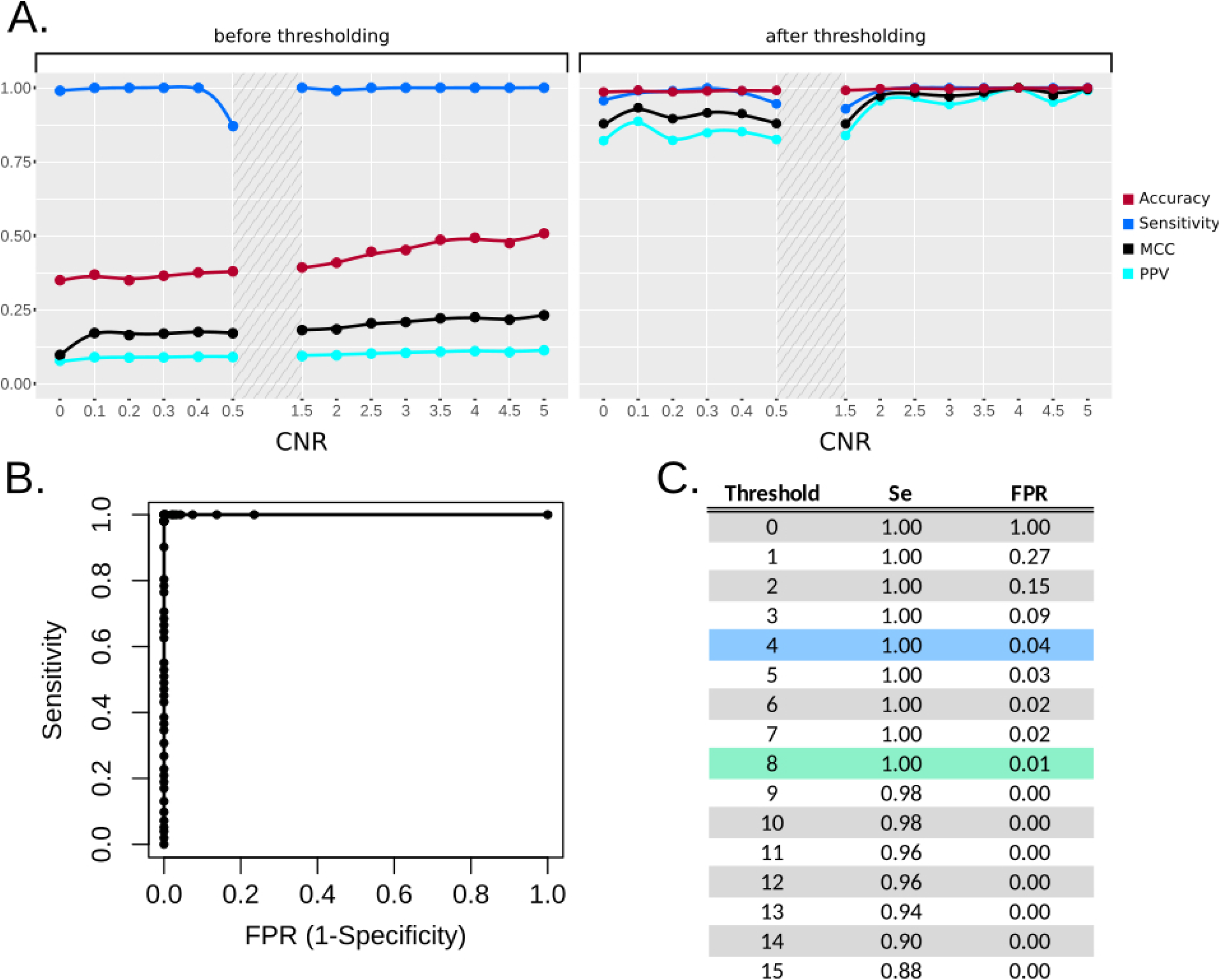
Performance assessment of the localisation score thresholding. A. Classification indicators for the detection of altered targets before (left panel) and after (right panel) thresholding. B. ROC curve. C. Associated table. FPR: false positive rate; Se: sensitivity; ROC: receiver operating characteristic.

As figure 3 shows, before thresholding the Se was close to 1 and the PPV and MCC were low (around 0.1 and 0.2, respectively). Acc was lower using the grouped targets (around 0.4), because, by construction, the total number of regions in the dataset is lower than the number of targets and therefore the number of TN is smaller. After thresholding, we observed that the PPV, MCC and Acc increased to reach values very close to 1 for CNRs over 2, while the Se had only slightly decreased, meaning that the score thresholding enables the TPs to be kept while the FPs are discarded. The localisation score thresholding approach can therefore be validated and represents an important improvement in the performance of our algorithm.

To characterise the dependence on the score, we iterated the threshold from 0 to 15, calculated the corresponding TPR and FPR and plotted a ROC curve (figure 4B). The corresponding table in figure 4C shows that, on the simulated data, CNVs with a score over 4 were 100% TPs and less than 5% FPs and CNVs with a score over 8 were 100% TPs and ~1% FPs.

### Evaluation of ifCNV on patient datasets

We then tested ifCNV on three real datasets. First, we used the ICR96 dataset and then used two in-house datasets for which we had NGS and aCGH results to compare to. Table 2 and Table 3 present the results.

**Table 2.** Classification indicators for ifCNV and the tools described in Morena-Cabrera et al. (ref) on the ICR96 dataset. Acc: accuracy; FN: false negative; FP: false positive; Sp: specificity; TN: true negative; TP: true positive.

| **Tool** | **TP** | **TN** | **FP** | **FN** | **Total** | **FNR** | **FPR** | **Se** | **Sp** | **PPV** | **Acc** | **MCC** |
| --- | --- | --- | --- | --- | --- | --- | --- | --- | --- | --- | --- | --- |
| ifCNV – auto | 296 | 27017 | 1858 | 0 | 28875 | 0 | 0.065 | 1 | 0.9350 | 0.1374 | 0.9363 | 0.3586 |
| ifCNV – None | 252 | 28875 | 0 | 44 | 28875 | 0.1486 | 0 | 0.8513 | 1 | 1 | 0.9984 | 0.9219 |
| DECoN | 286 | 28473 | 106 | 10 | 28875 | 0.0338 | 0.0037 | 0.9662 | 0.9963 | 0.7296 | 0.9959 | 0.8377 |
| panelcn.MOPS | 284 | 28236 | 343 | 12 | 28875 | 0.0405 | 0.012 | 0.9595 | 0.988 | 0.453 | 0.9877 | 0.6547 |
| CoNVaDING | 283 | 28068 | 511 | 13 | 28875 | 0.0439 | 0.0179 | 0.9561 | 0.9821 | 0.3564 | 0.9818 | 0.5778 |
| exomedepth | 283 | 28507 | 72 | 13 | 28875 | 0.0439 | 0.0025 | 0.9561 | 0.9975 | 0.7972 | 0.9970 | 0.8716 |
| CODEX2 | 275 | 28503 | 76 | 21 | 28875 | 0.0709 | 0.0027 | 0.9291 | 0.9973 | 0.7835 | 0.9966 | 0.8515 |

**Table 3.**
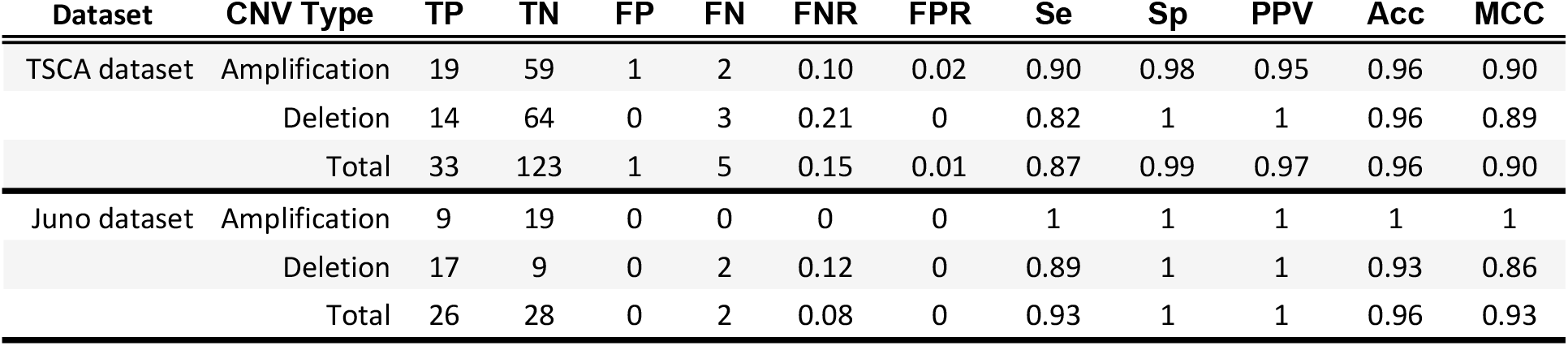
Classification indicators on the TSCA and the Juno datasets.

#### ICR96 dataset

To compare the performance of ifCNV to that of other methods,^18,24–26,34^ we compared our results to those obtained by Moreno-Cabrera et al.^35^ who benchmarked five widely used tools with the ICR96 dataset.^32^ This dataset, however, only possess one target per exon, rendering the advantage of the ifCNV score thresholding strategy not practicable in this case. As an alternative strategy, we decided to take advantage of the ability to change the *contamination* parameter of the IF. This main actionable parameter is the proportion of outliers in the dataset. By default, it can be set to a value between 0 and 0.5 and to ‘auto’, for an automatic detection of the proportion of outliers. In ifCNV we added the ability to set this parameter to ‘None’; it is then calculated as 1 on the number of samples in the dataset. We thus iterated several values of the *contamination* parameter and calculated the correspondent binary classification indicators (figure 5). We also compiled the results obtained with the two pre-set *contamination* values (‘auto’ and ‘None’) with the results from the other tools (Table 2). We observed that ifCNV exhibits performances in the same order of magnitude as the other tools, with the clear advantage of having an easily tunable parameter allowing the user to expect either no FNs (*contamination*=‘auto’) or no FPs (*contamination*=‘None’).

**Figure 5.**
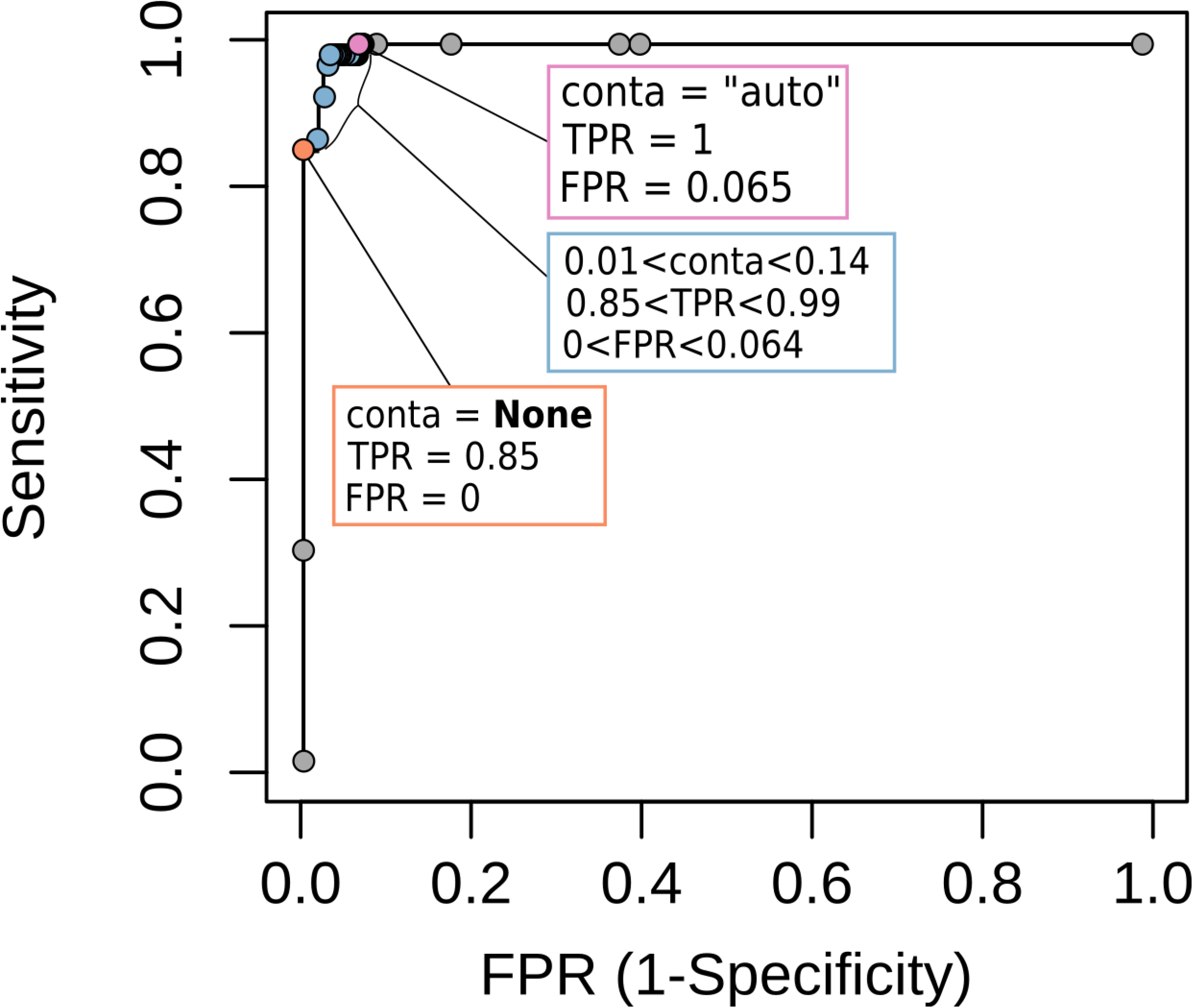
ROC curve for the ICR96 dataset. Some points of interest are highlighted: contamination = “auto” (pink), 0.01 < contamination < 0.14 (blue) and contamination = “None” (orange). TPR: true positive rate.

#### TSCA dataset

Next, we aimed to validate the performance of ifCNV on an in-house dataset (Table 3). Its particularities are: (i) it is composed of tumour samples containing variable percentages of altered cells, and (ii) it possesses a small number of targets per region (range: 1 to 40, median: 4). Using this dataset, we found that ifCNV correctly labelled 19 of the 21 amplifications present in the dataset; the 2 FN were measured as a gain of two copies (CNR=2) by aCGH. In addition, 14 of 17 deletions were detected with no FPs. The 3 undetected deletions were from samples that have a lower percentage of tumour cells.

#### Juno dataset

Finally, we also assessed our tool on a distinct library preparation approach with a larger number of targets per region composed of a larger panel than the TSCA dataset (range: 1 to 164, median: 17) from tumour samples (Table 3). Interestingly, we could detect all amplifications with no FPs and 17 out of the 19 deletions, leading to an overall Acc of 0.96 and MCC of 0.93. Interestingly, the 2 missed deletions are on a gene that represents only 0.4% of the panel (MTR=0.004).

## Discussion

In recent years, numerous computational methods for detecting and measuring CNVs from NGS data have been developed. However, most of these are based on the use of internal or external reference samples. To date, only a few have taken advantage of easy-to-use machine learning packages.^16,17,27^ Here, we describe ifCNV, a bioinformatics tool using a machine learning-based algorithm, allowing detection of CNVs without the need for a reference sample.

Moreover, in routine clinical practice, the variety of pathologies involving specific molecular alterations leads to a broad diversity in the datasets generated. Thus, in general, most CNV software that is developed for a specific data type has sub-optimal reliability for use in routine practice with diversified samples. In addition, most of the genetic workflows are either developed in python or R and, to our knowledge, no existing CNV detection tool is available in both languages.

ifCNV is, making it more easily to implement in pre-existing pipelines. Also, by successfully creating its own normal reference inside each analysed NGS run, ifCNV frees itself from any batch effect inherent to tools using external references. It also avoids the need for reference samples that are copy number neutral to be sequenced in the same batch, which is an efficient but not cost-effective solution. Furthermore, its efficiency and simplicity make it possible to run on hardware with limited computing resources.

Using simulated data, we demonstrate that ifCNV is highly reliable and adapts to several relevant clinical situations including: (i) when one CNV^pos^ sample is present in a set of several CNV^neg^ samples, (ii) when multiple CNV^pos^ samples are present in a set of several CNV^neg^ samples, (iii) when only one target is altered and (iv) when the CNRs are close to one, which can correspond to small alterations or to mixtures of normal and altered cells. ifCNV also performed well using datasets generated from amplicon- or capture-based libraries prepared from germline or somatic clinical samples.

Analyses of real data also demonstrated that ifCNV’s performance was comparable to that of other widely used tools,^35^ but with substantial specific advantages. Our solution has a tuneable control of the FPR thanks to localisation score thresholding and to the *contamination* parameter, which can both be optimised according to the dataset, by an entry-level user. ifCNV was also able to accurately detect CNVs in difficult samples, such as those composed of a mixture of normal cells and tumoral cells, which dilutes the CNR of samples.

In conclusion, ifCNV is a highly flexible tool that can detect CNVs in several types of clinical samples. ifCNV now represents an essential component of the cancer diagnosis pipeline that we routinely use to analyse samples from patients in our laboratory. We believe that the flexibility, high accuracy, easy implementation and low hardware infrastructure afforded by our method will help other laboratories in increasing their throughput and improve disease characterisation by accurate CNV detection.

